# Assessment of Robustness to Temperature in a Negative Feedback Loop and a Feedforward Loop

**DOI:** 10.1101/774042

**Authors:** Abhilash Patel, Richard M. Murray, Shaunak Sen

## Abstract

Robustness to temperature variation is an important specification for biomolecular circuit design. While cancellation of parametric temperature dependences has been shown to improve temperature robustness of period in a synthetic oscillator design, the performance of other biomolecular circuit designs in different temperature conditions is relatively unclear. Using a combination of experimental measurements and mathematical models, we assess the temperature robustness of two biomolecular circuit motifs — a negative feedback loop and a feedforward loop. We find that the measured responses in both circuits can change with temperature, both in the amplitude and in the transient response. We find that, in addition to the cancellation of parametric temperature dependencies, certain parameter regimes can also facilitate temperature robustness for the negative feedback loop, although at a performance cost. We discuss these parameter regimes of operation in the context of the measured data for the negative feedback loop. These results should help develop a framework for assessing and designing temperature robustness in biomolecular circuits.

## Introduction

An important specification for a design is that it function robustly in different environmental conditions. Temperature is a global environmental variable that can affect performance in multiple design contexts. Robustness to temperature changes is an important specification in contexts such as semi-conductor electronics [1] and instrumentation [2]. This temperature dependence arises because the underlying components, such as current-voltage characteristic in semiconductors, are temperature dependent. A standard way to engineer temperature robustness is to design and configure component temperature dependencies such that their cumulative effect cancels out. Because the properties of biological design substrates such as DNA, RNA, and proteins can also be temperature dependent, robustness to temperature changes is an important consideration for design in biology.

Robustness of biomolecular circuit function to temperature variations is an important problem in natural and synthetic contexts (Fig. 1). For example, in circadian rhythms, the mechanisms underlying the independence of period to temperature changes have been investigated [3, 4]. As another example, robustness of bacterial chemotaxis to temperature variations has also been investigated [5]. In both these contexts, the common principle for temperature robustness is to have “matched” parametric temperature dependencies, meaning that the temperature dependence of circuit parameters is such that their cumulative temperature dependence cancels out. Synthetic oscillator designs can also have temperature dependent periods [6], and relatively recently, it was shown how a temperature sensitive mutant transcription factor can be used to compensate for the effect of temperature [7]. We have investigated, primarily computationally, the propagation of temperature dependence in simple biomolecular circuit models, noting that in addition to matching temperature dependence of parameters, certain parameter regimes can also give temperature robustness [8, 9]. More recently, a multiscale approach was used to explain the effects of temperature in simple negative and positive feedback circuits in yeast [10]. These results provide some case studies highlighting principles and frameworks that can facilitate temperature robustness in biomolecular circuits.

**Figure 1:**
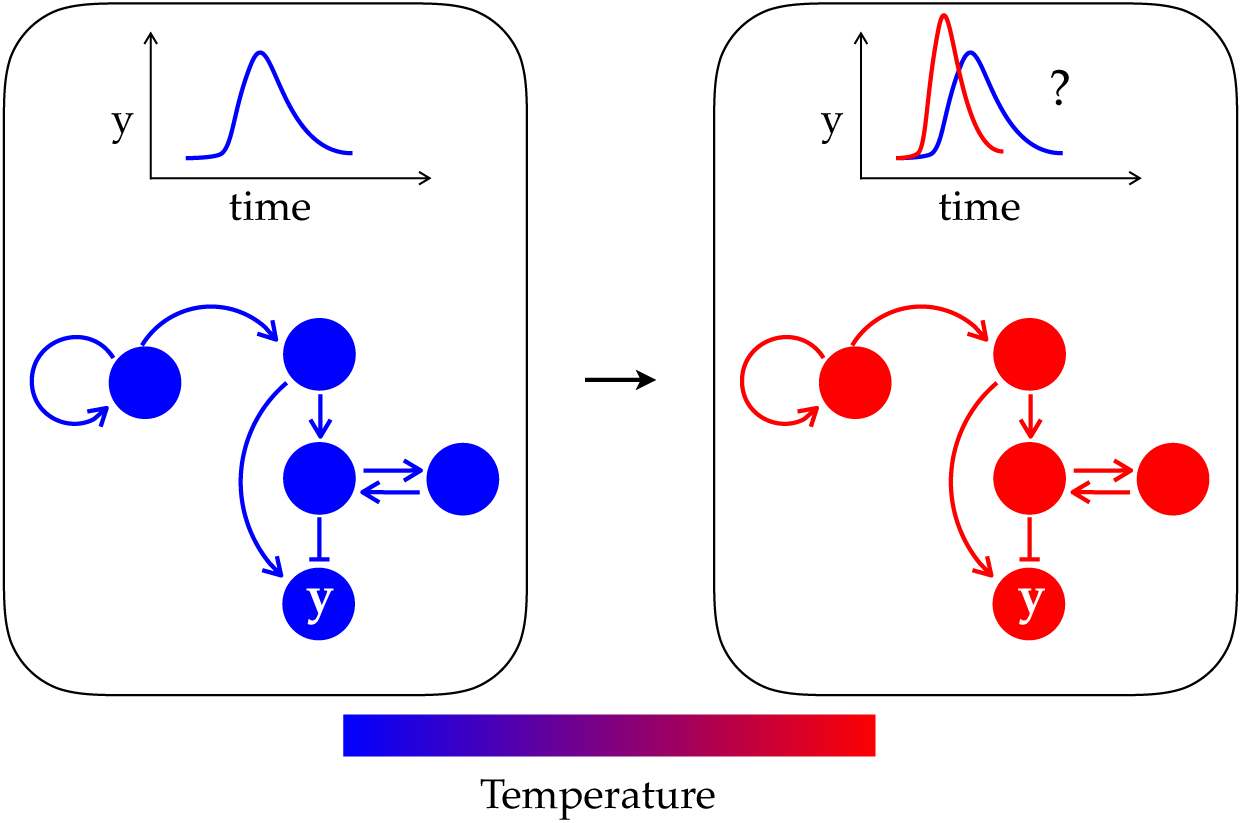
Biomolecular circuit response may change with temperature. Arrows represent interaction between the circle-shaped biomolecules. Blue and red represent cold and hot temperatures, respectively.

There are three notable aspects relating to temperature robustness in biomolecular circuits. First is the similarity in achieving temperature robustness between biomolecular contexts and other engineering contexts, through a suitable matching of temperature dependencies of the parameters. Second is the relative ease with which biomolecular circuit outputs can be measured in comparison with the biomolecular circuit parameters, which makes the design of such matching temperature dependencies a matter of finding the right mutant, if it exists. Third is the possibility that some parameter regimes of behaviour can facilitate temperature robustness. A characterisation of temperature robustness in standard biomolecular circuit designs may help in designing temperature robustness.

Here we ask about the extent of robustness to temperature changes in two biomolecular circuit motifs — a negative feedback loop and a feedforward loop. To address this, we use a combination of experimental measurements and mathematical models. We find that the response of these circuits changes with temperature, both in the amplitude and in the transient response. We analysed the underlying mathematical models, finding that in addition to having parameters with matched temperature dependencies, certain parameter regimes can also facilitate temperature robustness, although at a performance cost. For the negative feedback circuit, in particular, strong negative feedback can be more robust to temperature changes than weak negative feedback, but expression levels are lower. These results help develop a framework to assess and design robustness to temperature in biomolecular circuits.

## Results and Discussion

### Experimental Assessment of Temperature Dependence

We assessed the temperature dependence of two circuits — a previously investigated negative feedback loop [11] and a previously constructed feedforward loop realization (Addgene plasmid #45789). These two circuits belong to the set of circuits identified as widely occurring in genetic networks [12].

The negative feedback circuit has the transcriptional repressor TetR expressed under a *P*_*tet*_ promoter (Fig. 2a). TetR represses the *P*_*tet*_ reporter forming the negative feedback loop. The TetR protein is fused to Green Fluorescent Protein (GFP). The inducer anhydrotetracycline (aTc) can be used to tune the strength of the negative feedback through its inhibitory effect on the transcriptional activity of TetR.

**Figure 2:**
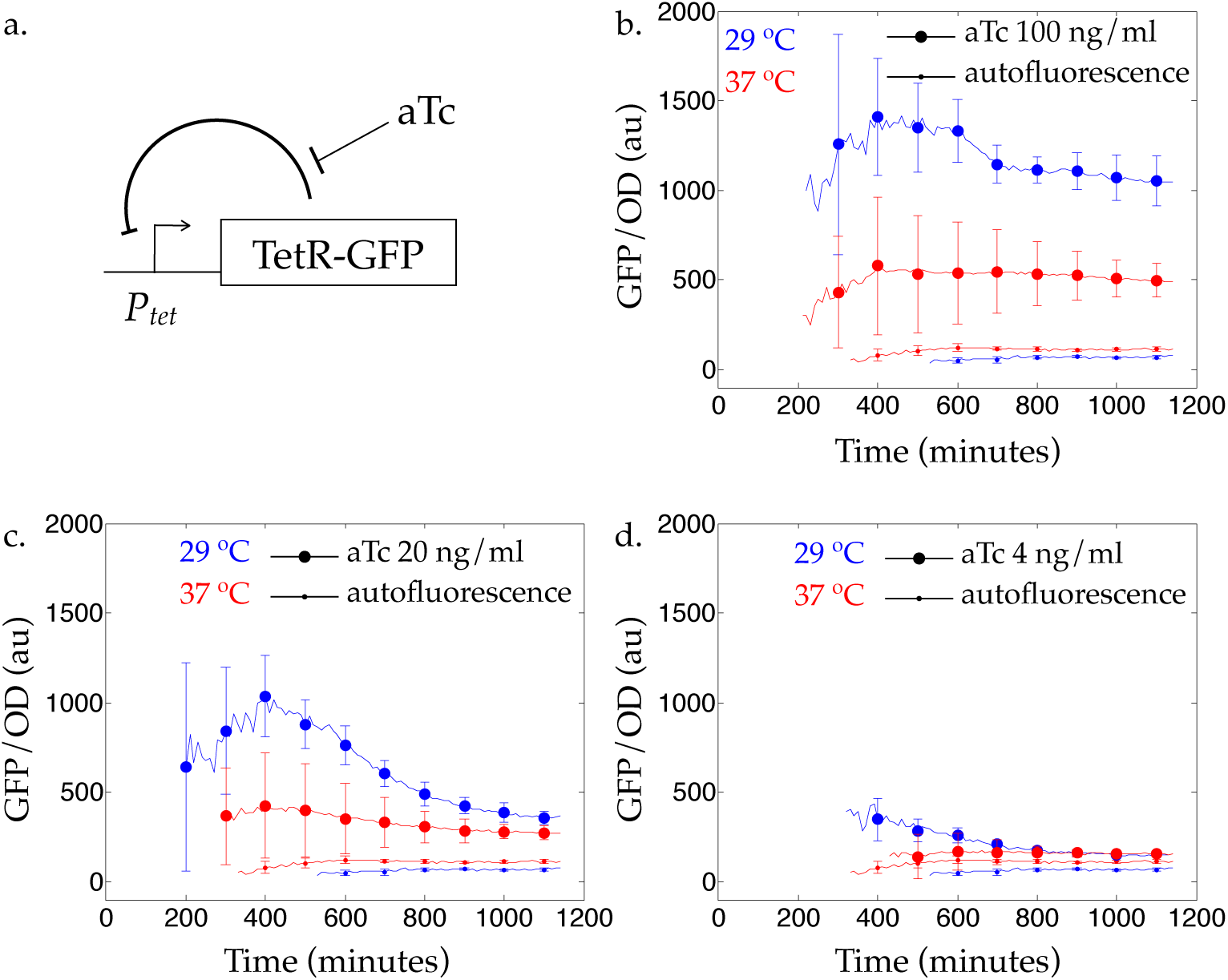
Temperature dependence of a negative feedback loop. a. Schematic of the circuit. b. Blue and red lines are responses measured at 29 °C and 37 °C, respectively. Circles and dots indicate the circuit response and autofluorescence, respectively. aTc concentration is 100 ng/ml. c. Same as in b. with aTc concentration 20 ng/ml. d. Same as in b. with aTc concentration 4 ng/ml.

We measured the response of the circuit at 29 °C and 37 °C(Fig. 2b–d; Methods). These measurements were at different concentrations of aTc. We find that the optical density-normalized fluorescence changes with temperature, both in the amplitude and the response. We note that temperature is a global variable and can affect other aspects of the measurement such as the GFP fluorescence and its dynamics as well as the aTc binding properties and its half-life.

The feedforward loop circuit consists of a transcriptional activator AraC, a transcriptional repressor TetR, and a degradation-tagged Green Fluorescent Protein GFP-ssrA (Fig. 3a.; Methods). AraC is constitutively expressed and is a transcriptional activator in the presence of the inducer arabinose. TetR is expressed from *P*_*BAD*_, an AraC-activated promoter. GFP-ssrA is expressed from a *P*_*BAD*_ promoter modified to have TetR operator sites so that it is repressible by TetR.

**Figure 3:**
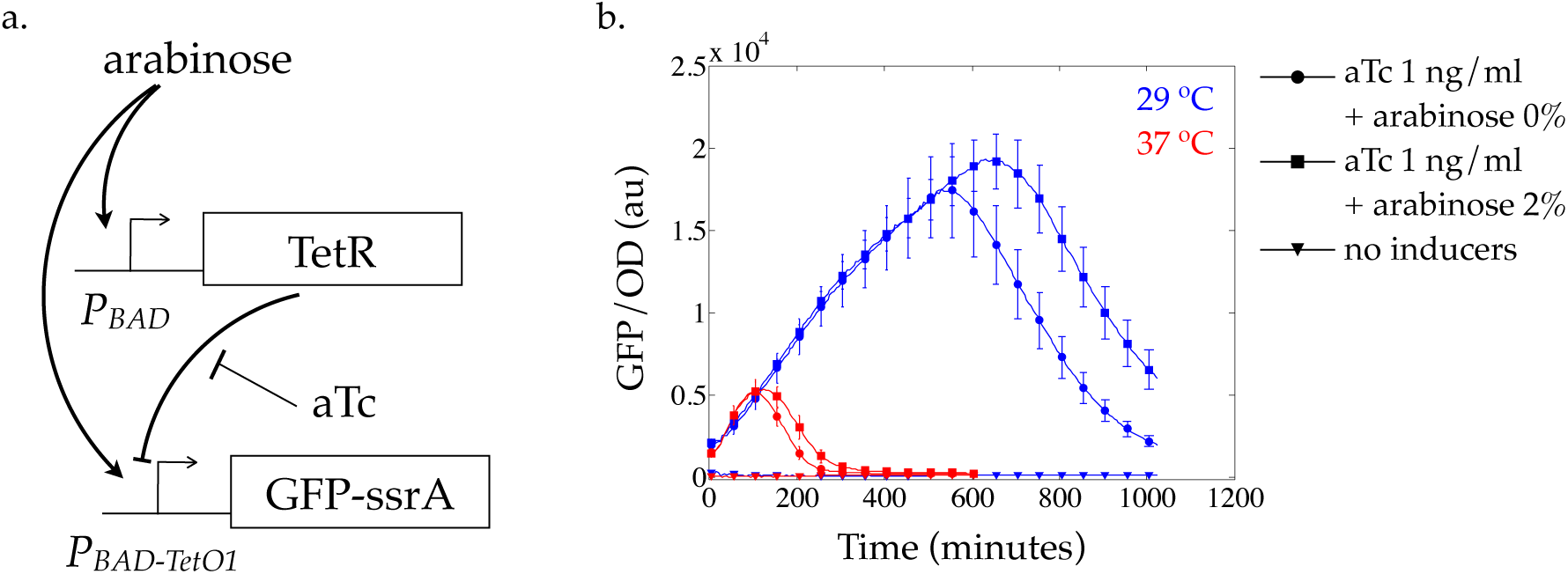
Temperature dependence of a feedforward loop. a. Schematic of the circuit. b. Blue and red lines are responses measured at 29 °C and 37 °C, respectively. Circles, squares, and triangles are responses for indicated inducer conditions.

We measured the response of the circuit at 29 °C and 37 °C (Fig. 3b.;Methods). We find that the optical density-normalized fluorescence shows a pulse response at both temperatures. While the circuit is designed to generate a pulse in GFP-ssrA expression in response to the addition of arabinose, a fixed level of aTc is required for the pulse to be observed, perhaps due to the dominant effect of TetR. In the presence of aTc, addition of arabinose has a small effect on increasing the amplitude of the pulse. The properties of the observed pulse response differ at these temperatures, which is apparent both in the amplitude of the pulse as well as in the time of the peak response.

These experimental measurements show that the overall response of two common circuits depends on temperature.

### Computational Assessment of Temperature Dependence

To complement the above experimental assessment, we considered simple models of these circuits to assess how the circuit response may depend on temperature.

For the negative feedback circuit, we considered a standard model of a protein *X* that transcriptionally represses its own expression and is degraded in a first-order process [11],

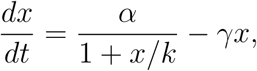

where *x* is the protein level, *α* is the maximal production rate, *k* is the DNA binding constant of X to its own promoter, and *γ* is the degradation rate constant. Nominal values of the parameters were as follows: *α* = 100 nM/hr, *k* = 10 nM, and *γ* = 1 /hr. As reference, we also considered a model without feedback,

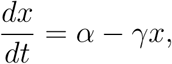

which can be exactly solved, 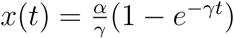 for a zero initial value of *x*.

To assess the temperature dependence of the response, both at steady-state and transiently, we assumed temperature dependencies of the parameters and computed the relative change in the response. We model the parametric temperature dependencies based on the temperature coefficient *Q*_10_, the amount that parameters are scaled when the temperature increases by 10 °C. As typical values of *Q*_10_ are in the range 2–3 [4], we assumed the *Q*_10_’s of *α* and *γ* to lie in this range. For *k*, which is itself a ratio of two rates, we assumed that the *Q*_10_ is in the range 0.66 − 1.5, which is the maximal range assuming that the numerator and denominator in the ratio vary independently. For a random choice of parameters around the nominal point (*M* = 100) and a random choice of *Q*_10_’s (*N* = 100) for each parameter set, we scaled the parameters according to the *Q*_10_’s and computed the steady-states and the transient response (Fig. 4a, d). The histogram of the temperature coefficients of the steady-state, *Q*_*xss*_ are shown in Fig. 4b, e. For the negative feedback model, the mean is 1.03 and the standard deviation is 0.14. For the model without feedback, the mean is 1.01 and the standard deviation is 0.17. A *Q*_10_ of exactly one corresponds to perfect temperature robustness — there is no change with temperature. In both circuits, we observed that the mean is close to one even though the temperature coefficients of underlying parameters do not have a mean close to one. This temperature robustness effect corresponds to the situation when the temperature dependencies scale similarly and there is a net cancellation.

**Figure 4:**
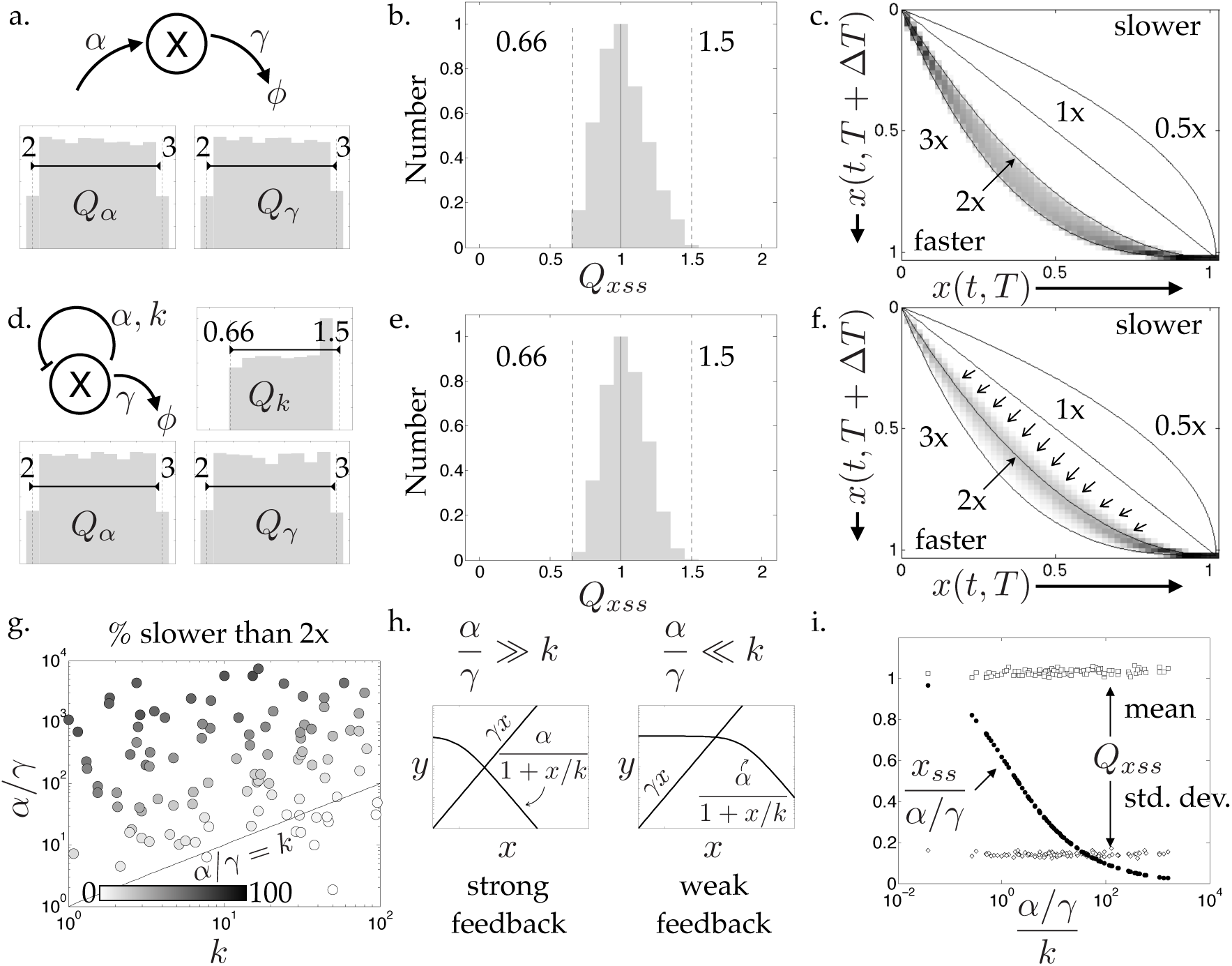
Computational assessment of temperature dependence of a negative feedback loop. a. Schematic of a circuit with no feedback. Histograms show assumed temperature coefficients of parameters. b. Histogram of the computed temperature coefficient of the steady-state. c. Normalised transient response after temperature shift is plotted versus the the initial transient response. Grayscale indicates the sum of all trajectories at a particular grid location. Solid lines represent curves corresponding to an exponential response and responses that are identical, faster, or slower. Same as in a., but for a negative feedback circuit. e. Same as in b., but for a negative feedback circuit. f. Same as in c., but for a negative feedback circuit. Arrows highlight the portion of the plot that is different from that in c. g. Dots are colour-coded based on the fraction of realisations that are less than two-fold faster as estimated by f. These are plotted in the space of the parameter combinations *α/γ* and *k*. Solid line is *α/γ* = *k*. h. Interpretation of a weak and a strong negative feedback regime based on relative values of parameter combinations *α/γ* and *k*. i. Black solid circles represent the steady-state value normalized by *α/γ* as a function of the negative feedback strength *α/γ/k*. White squares and diamonds represent the mean and standard deviations of the temperature coefficient of the steady-state as a function of the negative feedback strength *α/γ/k*, respectively.

To assess the transient response, we plotted the response, normalised to steady-state, after the parameters were scaled versus the response before scaling (binned versions are shown in Fig. 4c, f). If the responses were unchanged, this curve would lie on the indicated diagonal. As reference, exponential responses with time constants that are half, double, and triple are plotted. These correspond to a slowing down of the response by twofold, speeding up of the response by twofold, and a speeding up of the response by threefold, respectively. For the circuit without feedback, all responses lie in the region spanned by the twofold and threefold speed-up curves, corresponding exactly to a *Q*_10_ in the range of 2–3 for the parameter *γ* (Fig. 4c). For the negative feedback circuit, the responses lie on both sides of the two-fold speedup curve (Fig. 4f), suggesting that there are parameter regimes that facilitate robustness to temperature.

We further investigated this region of the plot. We found that for all parameters, there were some scalings which were less than twofold faster than the original response. When we counted the fraction of scalings, we found that a larger fraction occurred for parameters satisfying the relation *α/γ > k* (Fig. 4g). As *α/γ* is the maximum level of *x*, this corresponds to a region in parameter space where negative feedback interaction is dominant and *α/*(1 + *x/k*) ≈ *α/x/k* (Fig. 4h). This is the region that enhances robustness to temperature. We checked whether this region affects the temperature dependence of the steady-state (Fig. 4i). The mean and standard deviations of the steady-state temperature coefficients did not change as a function of the parameter *α/γk*. However, the absolute value of the steady-state relative to the maximum allowable value *α/γ* decreases as *α/γk* increased. This suggests that there could be a performance cost to temperature robustness, in terms of a lower steady-state value, for circuits operating in the strong negative feedback regime.

We repeated this analysis for the incoherent feedforward loop circuit (Fig. 5). We considered the standard model [13],

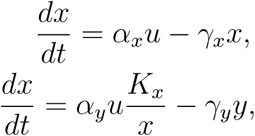

where *u* is the input, *y* is the output protein level, and *x* is the intermediatory protein that, like *y*, is activated by *u*, but which represses *x*. The parameters *α*_*x*_ and *α*_*y*_ represent the production rates of *x* and *y* respectively. Similarly, the parameters *γ*_*x*_ and *γ*_*y*_ represent the degradation rate constants of *x* and *y* respectively. *K*_*x*_ is the DNA binding constant of *X* to the promoter of *Y* and uses the approximation that the repression function *K*_*x*_*/*(*x* + *K*_*x*_) ≈ *K*_*x*_*/x*. Nominal values of the parameters were: *α*_*x*_ = 100 nM/hr, *K*_*x*_ = 10 nM, *γ*_*x*_ = 1 /hr, *α*_*y*_ = 100 nM/hr, *γ*_*y*_ = 1 /hr, and the input *u* : 1 → 2.

**Figure 5:**
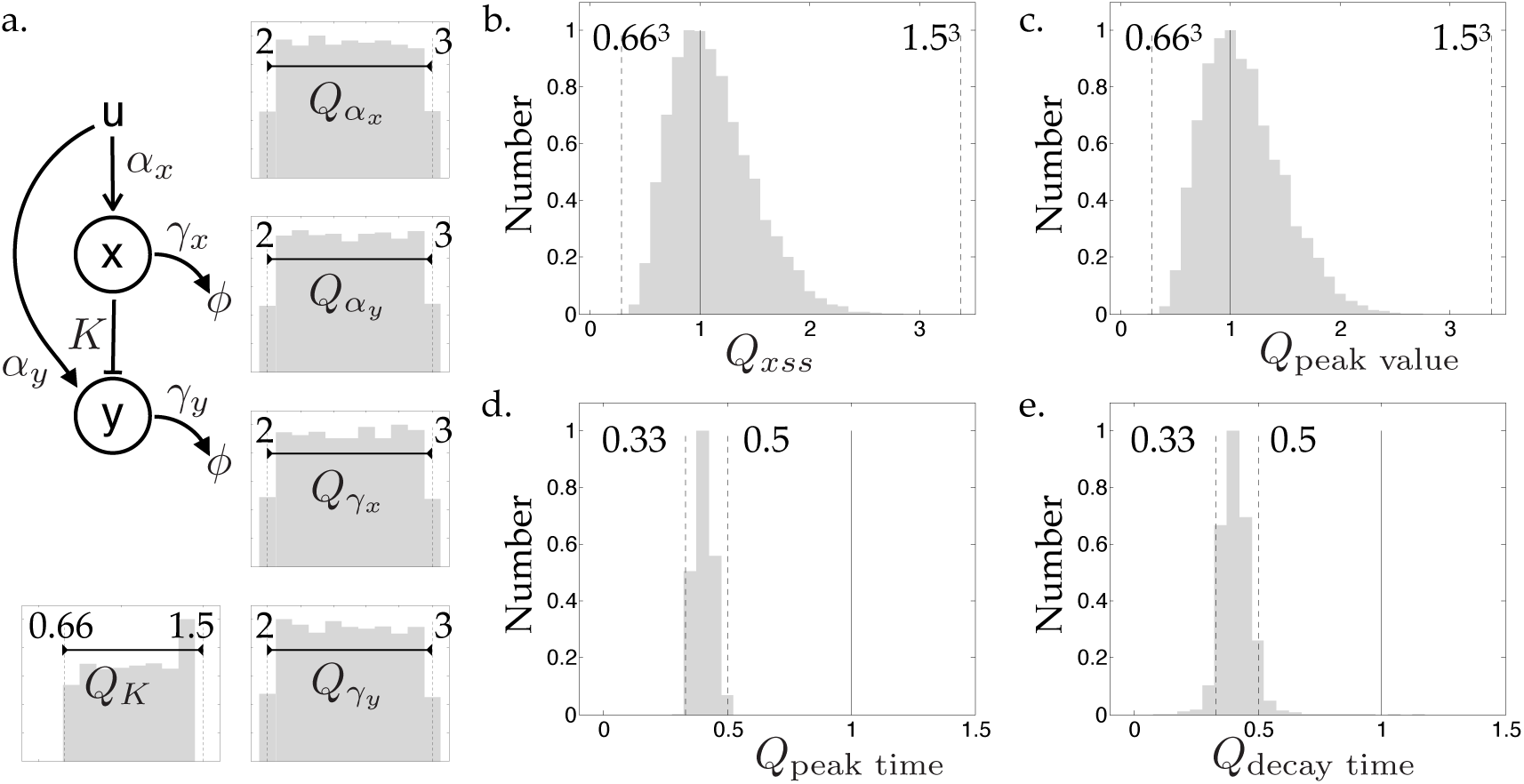
Computational assessment of temperature dependence of a feedforward loop. a. Schematic of a feedforward loop. Histograms show the assumed temperature coefficients of the parameters in the computations. Histograms of the computed temperature coefficient of b. steady-state, c. peak value of the transient response, d. peak time of the transient response, and e. decay time of the transient response.

We assessed the temperature dependence of the steady-state and transient response of *y* due to the temperature dependence of the parameters, modelled using the temperature coefficient *Q*_10_. The *Q*_10_’s of *α*_*x*_, *γ*_*x*_, *α*_*y*_, and *γ*_*y*_ were taken in the range 2–3 and the *Q*_10_ of *K*_*x*_ was taken in the range 0.66–1.5 (Fig. 5a.).

The distribution of the temperature coefficients of the steady-state has a mean of 1.11 and a standard deviation of 0.36 (Fig. 5b). As above, the mean temperature coefficient close to one indicates an overall temperature robustness effect. As the analytical steady-state expression of the feedforward loop circuit is *K*_*x*_*α*_*y*_*γ*_*x*_*/*(*α*_*x*_*γ*_*y*_), a mean close to one indicated that the parametric temperature dependencies scale similarly and there is an effective cancellation. The worst-case range of the *Q*_10_’s is expected to be (2*/*3)^3^, (3*/*2)^3^. Generally, if the sets of parameters *α*_*x*_, *α*_*y*_ and *γ*_*x*_, *γ*_*y*_ have matching temperature dependences, due to the similar nature of processes lumped as these parameters, then we may get a temperature compensatory effect.

We assessed the temperature dependence of transient response by considering three metrics — the peak amplitude, the time to rise to the peak amplitude, and the time to decay from the peak amplitude to 5% of its final value. The histogram of the temperature coefficients of each of these are shown in Fig. 5c–e. The peak amplitude has a largely similar histogram as that of the steady state, with a mean of 1.12 and a standard deviation of 0.36. The histograms of the rise time and decay time have a mean of 0.40 and 0.41, respectively. Both these histograms fall in the range 1*/*3 and 1*/*2. This suggests the dominant effect of the degradation rate parameters, as the timescale properties, which vary as the inverse of the degradation rate parameters, have temperature coefficients in the range 1*/*3–1*/*2.

### Discussion of the experimental measurements in the context of computations

The computations for the negative feedback circuit suggest that, in addition to matched temperature dependencies, certain parameter regimes can also facilitate temperature robustness, at a performance cost, similar to what can be observed in a two-state model [8]. Experimental measurements seem consistent with this (Fig. 2), as the change in the trajectories as temperature is increased from 29 °C to 37 °C is smaller for high aTc (strong feedback) in comparison with low aTc (weak feedback) and the expression levels are lower for strong fedback. However, two aspects of the experimental conditions are not in the numerical computations. First aspect is the possible temperature dependence of the GFP properties such as its brightness and maturation dynamics, which may be different for different temperatures. Second aspect is the use of aTc levels to access different parameter regimes as the binding of aTc to TetR may also depend on temperature, implying a comparison of circuits with different effective DNA binding affinities of TetR. To address the possible change in the brightness of GFP with temperature, we performed a melt curve of GFP inside cells containing the negative feedback circuit, finding that its fluorescence decreases by 9.39 *±* 2.87% (N = 2) from 30 °C to 37 °C (Supplementary Fig. 5). This suggests that, as far as brightness of this GFP is concerned, the trajectories for different temperatures may be compared with a small error.

We had previously reported the effect of temperature on a similar negative feedback loop in a cell-free context [14]. Based on the analysis presented here, we revisited this data. The data is comprised of the response of the negative feedback circuit, at two different aTc levels corresponding to strong and weak feedback, and a constitutive promoter in a cell-free context measured at four different temperatures — 26 °C, 29 °C, 33 °C, and 37 °C. (Fig. 6a–c; see the Methods section). We plotted the time it takes to reach half the final value, expecting that the strong negative feedback would minimize the change in this quantity with temperature. This is observed (Fig. 6d), both when comparing strong feedback to weak feedback and when comparing strong feedback to the constitutive promoter, providing experimental evidence for the temperature robustness effect of negative feedback. Next, we plotted the trajectories for the strong negative feedback, weak negative feedback, and constitutive promoter against each other to compare with the computational expectation (Fig. 2c,f), expecting that the trajectories for the strong negative feedback would be closer to each other in comparison with the other two cases. We see evidence of this (Fig. 6e,f) when the responses from 26 °C to 33 °C or to 37 °C are compared. Overall, these provide experimental evidence for the temperature robustness effect of negative feedback.

**Figure 6:**
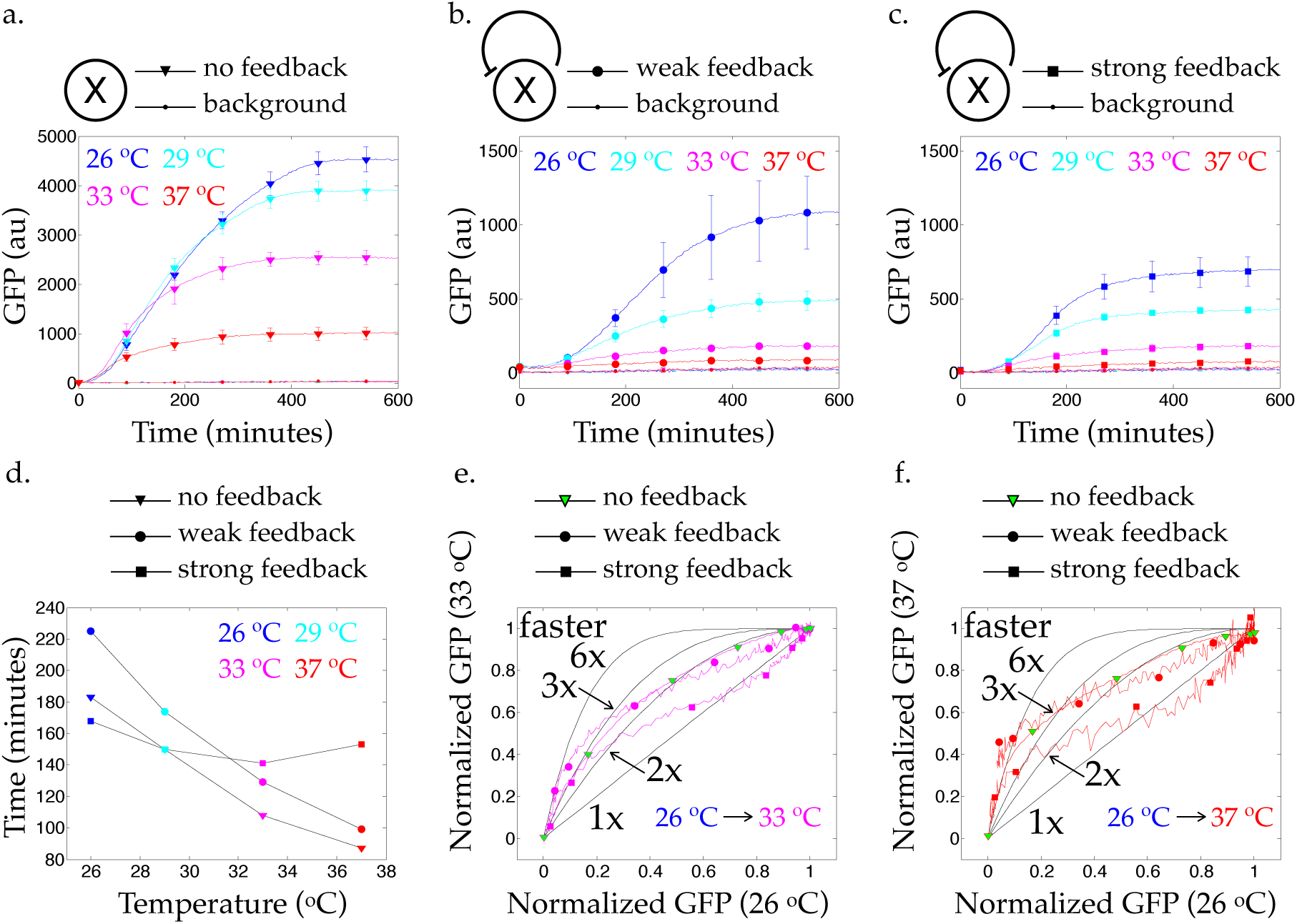
Temperature dependence of a negative feedback circuit in a cell-free context. Trajectories for a. a constitutive promoter, b. weak negative feedback loop, and c. strong negative feedback loop for 26 °C (blue line), 29 °C (cyan line), 33 °C (magenta line), and 37 °C (red line). Error bars represent standard deviations of three independent measurements. d. Time to reach half the final value for the above conditions. e. Response at 33 °C normalized by its final value versus the response at 26 °C normalized by its final values for the above conditions. Black lines represent guides for when an exponential response speeds up by the indicated numbers. Green symbol represents the no feedback case, for clarity. f. Same as above except the response on the y-axis is at 37 °C normalized by its final value.

We noted from the feedforward loop computations that the dominant temperature-dependent effect on the peak time is due to the degradation-related parameters (Fig. 5d). We find that this is consistent with the experimental measurements as the ≈ 72% decrease in peak time as temperature increases from 29 °C to 37 °C (Fig. 3b) correlates well with the ≈ 72% increase in the cell growth rate (assessed from the time to reach OD = 0.5, Supplementary Figure 4), which functions like degradation parameter in the feedforward model for the protein X.

## Summary

Assessing and designing robustness to temperature in biomolecular circuits is an important challenge for their possible application. Using a combination of experimental measurements and mathematical models, we address this using two biomolecular circuit motifs — negative feedback and feedforward loop. We find that the overall response of their circuit realizations changes with temperature, both in the amplitude and transient response. We analysed mathematical models of these circuits, highlighting that, in addition to having parameters with matched temperature dependencies, certain parameter regimes can also facilitate temperature robustness, albeit at a performance cost. We discuss the experimental measurements of the negative feedback loop in the context of the role of parameter regimes in facilitating temperature robustness. These results contribute to a framework for assessing and designing temperature robustness in biomolecular circuits.

## Materials and Methods

The plasmids and bacterial strains used as well as the methods for experimental measurements, data analysis, and computations are described below.

### Plasmids and Bacterial Strains

The negative feedback loop was obtained from Prof. M. B. Elowitz [11]. The plasmid is pZS*21tetR-egfp and is in the *E. coli* DH5*α*.

The feedforward loop pBEST-OR2-OR1-Pr-araC, pBAD-TetR, pBAD-TetO1-deGFP-ssrA is a plasmid from the lab of Prof. Richard M. Murray (Addgene plasmid # 45789; http://n2t.net/addgene:45789; RRID:Addgene 45789). This was transformed into the *E. coli* MG1655.

### Measurements

For the negative feedback loop, two strains, one containing the plasmid and the other without any plasmid, were grown overnight in minimal M9CA media (1x M9 salts, 1.0% glucose, 0.1% Casamino Acids, 0.5 *µ*g/ml Thiamine, 0.2 mM *MgSO*_4_, 0.1 mM *CaCl*_2_, pH 7.0, from Teknova) at required temperature (29 °C or 37 °C). The culture with the plasmid was grown in the presence of the kanamycin (50 ng/ml). These were diluted 1 : 100 in fresh media with the antibiotic. 190 *µ*l of each strain and of the media was placed in wells of a 96-well plate (Perkin Elmer). The negative feedback strain was also placed in other wells where following final concentrations of inducer aTc (in *µ*g/ml) were added to make total volume 200 *µ*l: 100, 20, 4. Each sample was placed in triplicate. The plate was incubated in a plate reader (Biotek Synergy H1) for 19 hours with shaking (double orbital, continuous, 237 cpm) and at the required temperature. Every 10 minutes, fluorescence (ex/em = 485/ 530 nm, gain 61) and optical density (absorbance at 600 nm) were measured. The measurements of the well containing only media provided the background for these measurements. These measurements were repeated on different days for 29 °C (N = 4 days) and 37 °C (N = 3 days). The feedforward loop strain was grown overnight in LB media supplemented with Ampicillin (100 *µ*g/ml). It was subsequently diluted in minimal media with the antibiotic and incubated with the inducer aTc (1 ng/ml) for two hours. Next, 0.2 % of the inducer Arabinose was added to the incubated culture and 200 *µ*l of it was placed in a 96 well plate (Eppendorf). The plate was placed in a plate reader (Biotek Synergy H1) and incubated for 10 hours at 37 °C and for 16 hours at 29 °C. Fluorescence (ex/em = 485/ 525 nm) and optical density (absorbance at 600nm) of each culture were measured every 5 minutes with a 2 minute shaking in between the readings. The samples were placed in duplicate and the above protocol was repeated for N = 3 days at each temperature.

### Data Analysis

All data analysis was performed in MATLAB.

For the negative feedback loop, the fluorescence and optical density of all samples across all days were used to get a sample mean and a sample standard deviation. The mean of the media background (fluorescence and optical density) was subtracted from the sample means. In these traces, there is an initial period where the background and samples overlap in the sense of the error bars given by the respective standard deviations. Therefore, for each sample, the time after which there is no overlap was considered. Neglecting the initial time measurements avoids the divide-by-zero (or a small number), which can result in large fluctuations in the optical density-normalized fluorescence. The optical density and fluorescence data are shown in Supplementary Fig. 1. The standard deviations of the optical density and fluorescence were obtained from the day-to-day samples. The optical density-normalized fluorescence values are shown in Supplementary Fig. 2. The standard deviation of the optical density-normalized fluorescence were obtained from these via quadrature assuming that they were uncorrelated. The data at 29 °C showed substantial variation relative to the data at 37 °C. One possibility could be that the temperature controller of the plate reader was more uniform at 37 °C (Suppplementary Fig. 3). As the temperature profiles of the first two repeats looked similar, these two readings were considered for the data at 29 °C shown in Fig. 2b-d.

For the feedforward loop, the background media was first subtracted and the mean and standard deviation are calculated as above. The optical density-normalized fluorescence values for 29 °C (N = 2 days) and 37 °C (N = 3 days) are shown in Fig. 3. The values for all three days as well as the optical density and fluorescence values are shown in Supplementary Fig. 4.

### Numerical Computations

Ordinary differential equations were solved in MATLAB using the function ode23s.

### Cell-free Experiments

These experiments were performed in a cell-free transcription translation system [15, 16]. The negative transcriptional feedback circuit used for the experiment contained a transcriptional repressor protein TetR expressed under a self-repressible promoter (Addgene plasmid # 45774; http://n2t.net/addgene:45774; RRID:Addgene 45774). TetR was fused to the green fluorescent protein variant deGFP and measured using excitation and emission at wavelengths 485 nm and 525 nm, respectively. The constitutive promoter is the lambda repressor Cro promoter (OR2-OR1-*P*_*r*_). It drives the expression of deGFP and is a positive control for the functioning of the cell-free reactions. These measurements were performed in a multilabel plate reader (BioTek Synergy H1) with measurement intervals set at 3 minutes for a total duration of 10 hours. These were then repeated at the different desired temperatures. The aTc concentration of 0.5 *µ*g/ml corresponded to relatively strong negative feedback and that of 5 *µ*g/ml corresponded to relatively weak negative feedback.

## Acknowledgements

We thank C. A. Hayes for her help with the cell-free experiments.

