## Supplementary Figures for "Assessment of Robustness to Temperature in a Negative Feedback Loop and a Feedforward Loop"

### Supplementary Figure 1.

a. Optical density and b. Fluorescence values of the data from the Negative Feedback circuit.

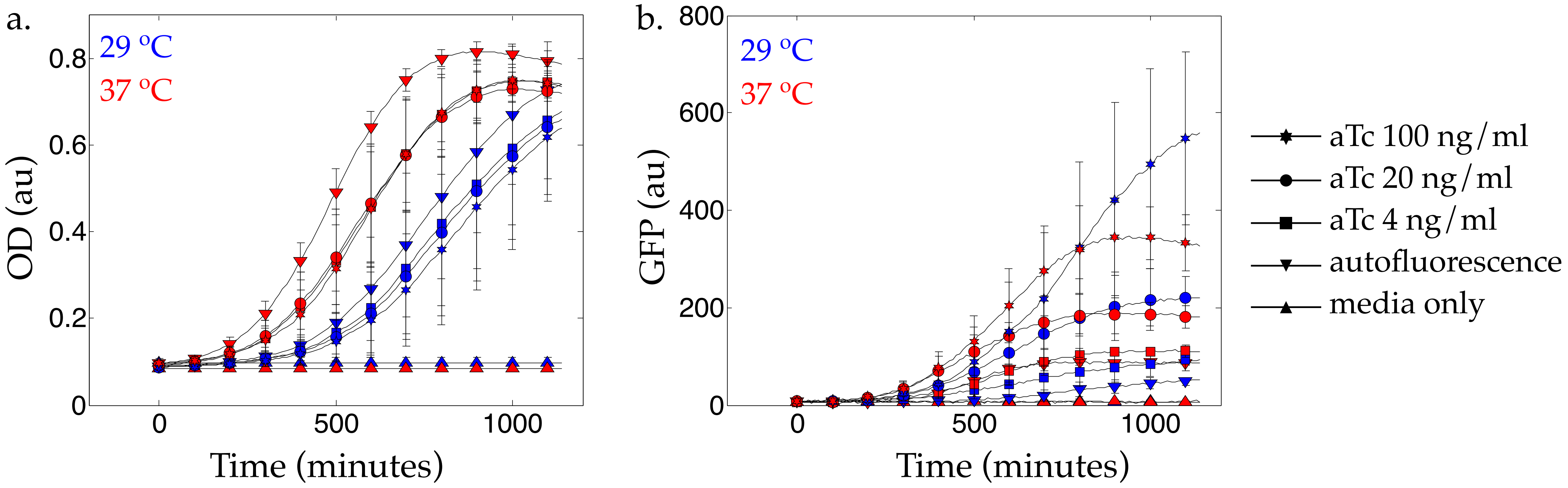

#### Supplementary Figure 2. Experimental assessment of negative feedback circuit.

Blue and red lines are the measured responses at 29 $^{o}$C and 37 $^{o}$C, respectively. Circles and dots indicate the response of the negative feedback loop and autofluorescence, respectively. aTc concentrations are indicated.

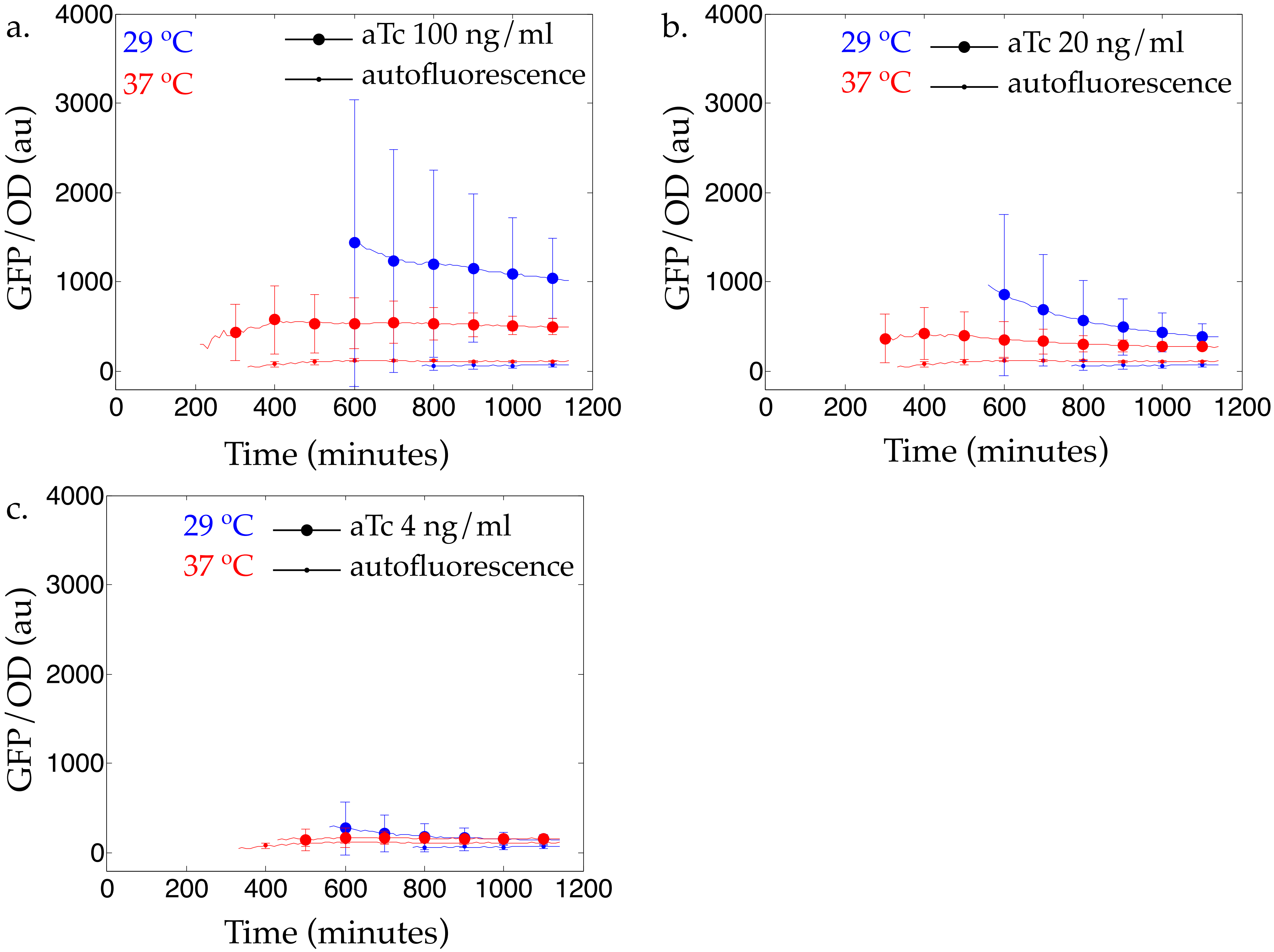

### Supplementary Figure 3.

Temperature profiles for the experimental assessment of negative feedback. Solid blue lines are the data that are plotted in Fig. 2.

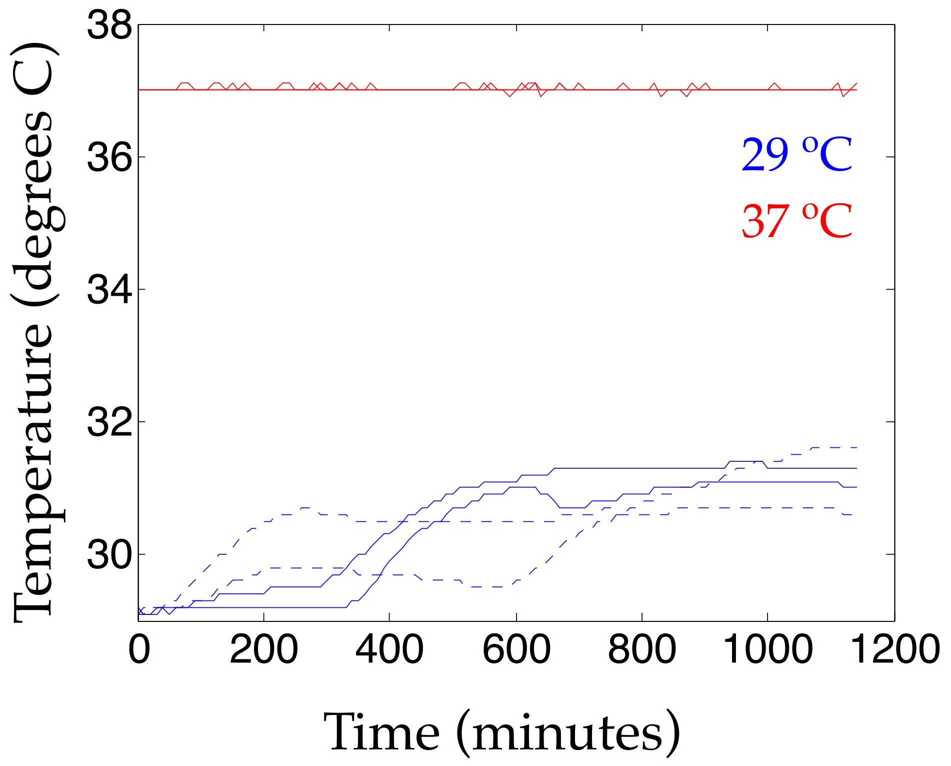

### Supplementary Figure 4.

a. Optical density, b. Fluorescence, and c. Optical density-normalized fluorescence values of the data from the Feedforward Loop Circuit.

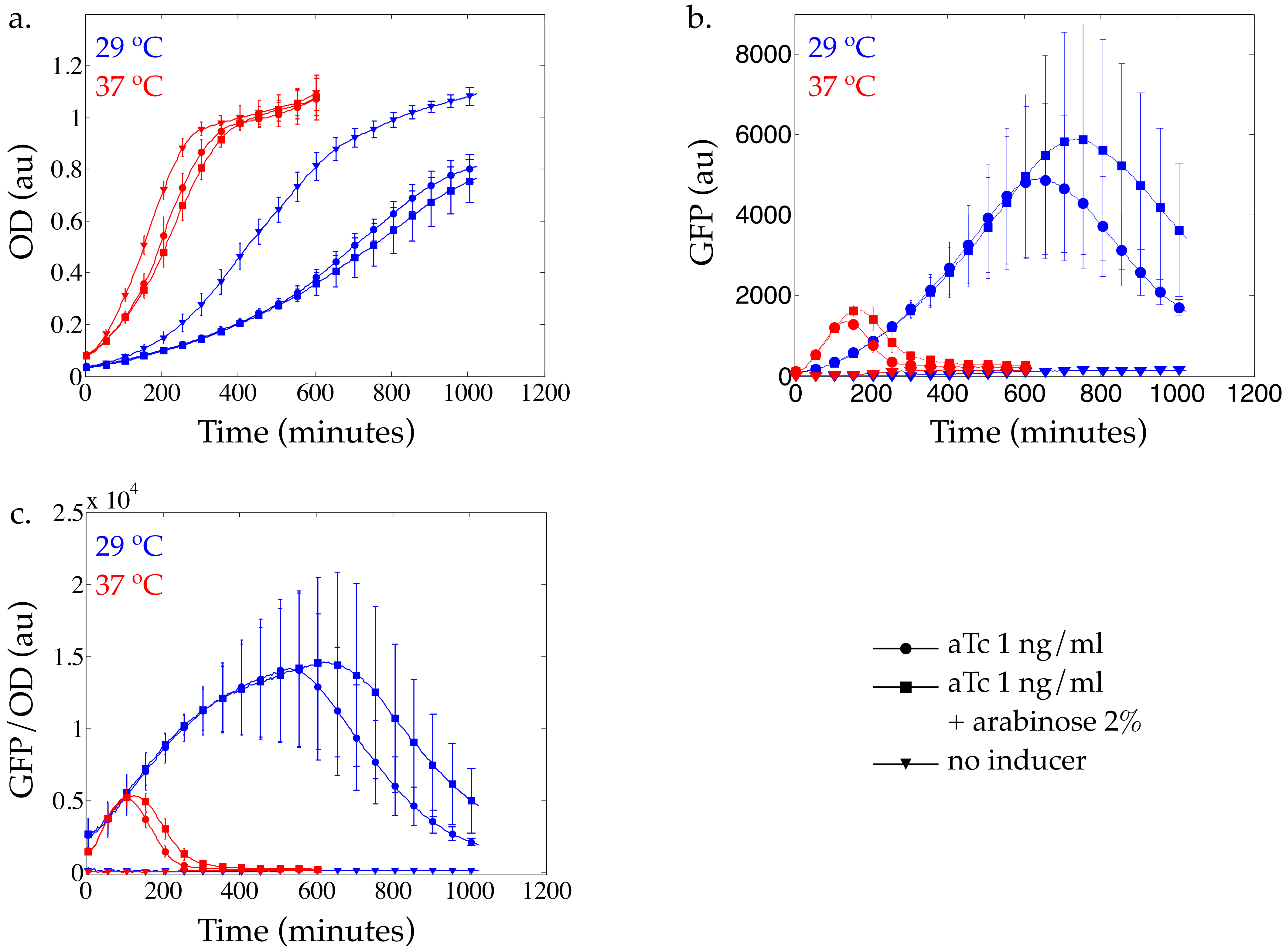

### Supplementary Figure 5.

**Melt curve of GFP inside cells containing negative feedback circuit** Cells were grown overnight in minimal media and antibiotics at 30 $^{o}$C. They were subsequently diluted 1:100 in fresh minimal media and antibiotics. 100 ng/ml aTc was added. This was incubated at 30 $^{o}$C for 24 hours. A melt curve was measured in a qPCR machine (Bio-Rad) from 25 $^{o}$C to 65 $^{o}$C in increments of 0.5 $^{o}$C and with 20 seconds at each temperature. Measurement was done via the SYBR channel. Measurements from a sample containing only media were subtracted.

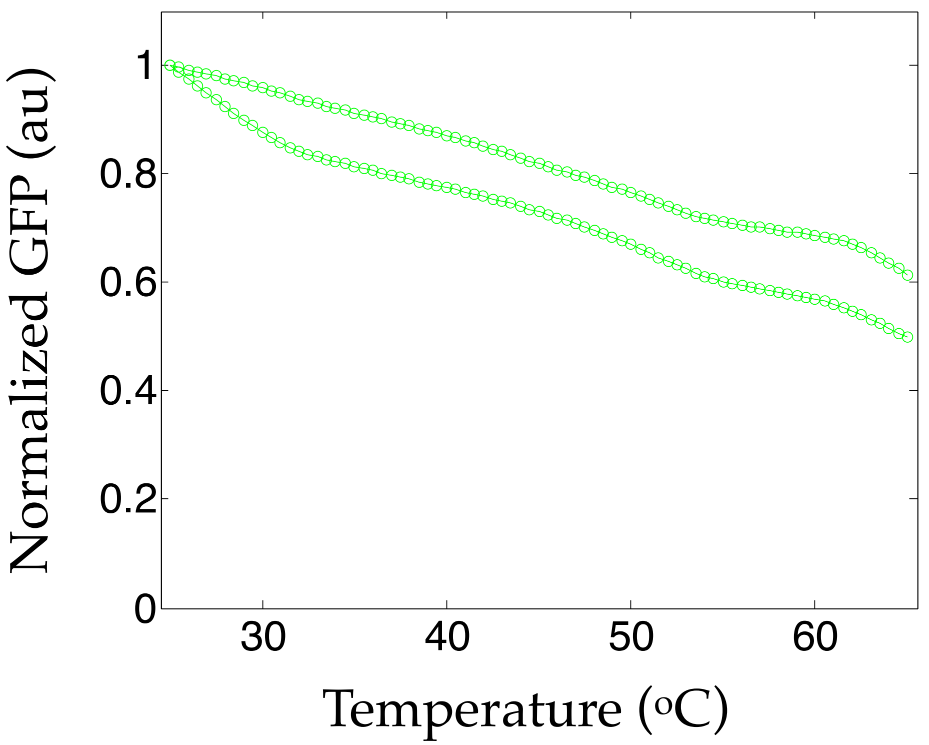
